# Neuronal circuits integrating visual motion information in *Drosophila*

**DOI:** 10.1101/2021.12.20.473513

**Authors:** Kazunori Shinomiya, Aljoscha Nern, Ian A. Meinertzhagen, Stephen M. Plaza, Michael B. Reiser

## Abstract

The detection of visual motion enables sophisticated animal navigation, and studies in flies have provided profound insights into the cellular and circuit basis of this neural computation. The fly’s directionally selective T4 and T5 neurons respectively encode ON and OFF motion. Their axons terminate in one of four retinotopic layers in the lobula plate, where each layer encodes one of four cardinal directions of motion. While the input circuitry of the directionally selective neurons has been studied in detail, the synaptic connectivity of circuits integrating T4/T5 motion signals is largely unknown. Here we report a 3D electron microscopy reconstruction, wherein we comprehensively identified T4/T5’s synaptic partners in the lobula plate, revealing a diverse set of new cell types and attributing new connectivity patterns to known cell types. Our reconstruction explains how the ON and OFF motion pathways converge. T4 and T5 cells that project to the same layer, connect to common synaptic partners symmetrically, that is with similar weights, and also comprise a core motif together with bilayer interneurons, detailing the circuit basis for computing motion opponency. We discovered pathways that likely encode new directions of motion by integrating vertical and horizontal motion signals from upstream T4/T5 neurons. Finally, we identify substantial projections into the lobula, extending the known motion pathways and suggesting that directionally selective signals shape feature detection there. The circuits we describe enrich the anatomical basis for experimental and computations analyses of motion vision and bring us closer to understanding complete sensory-motor pathways.

## Introduction

The *Drosophila melanogaster* visual system has been crucial for uncovering circuit mechanisms of many neural computations, such as detecting visual motion, looming, and color opponency^1–8^. Genetic driver lines enable functional studies of these computation^9–13^, often testing circuit hypotheses suggested by recent connectomes based on three-dimensional electron microscopy (3D-EM). The fly optic lobe has four major neuropils (lamina, medulla, lobula, and lobula plate; Figure 1A) that are characterized by columnar neurons connecting these structures, and striking layer patterns housing these connections. The diversity of optic lobe neuron types has been well documented using Golgi’s and silver staining methods^14,15^, and in recent years, genetic driver lines for cell-type-specific expression and new tools for neuroanatomy^11,16,17^.

**Figure 1.**
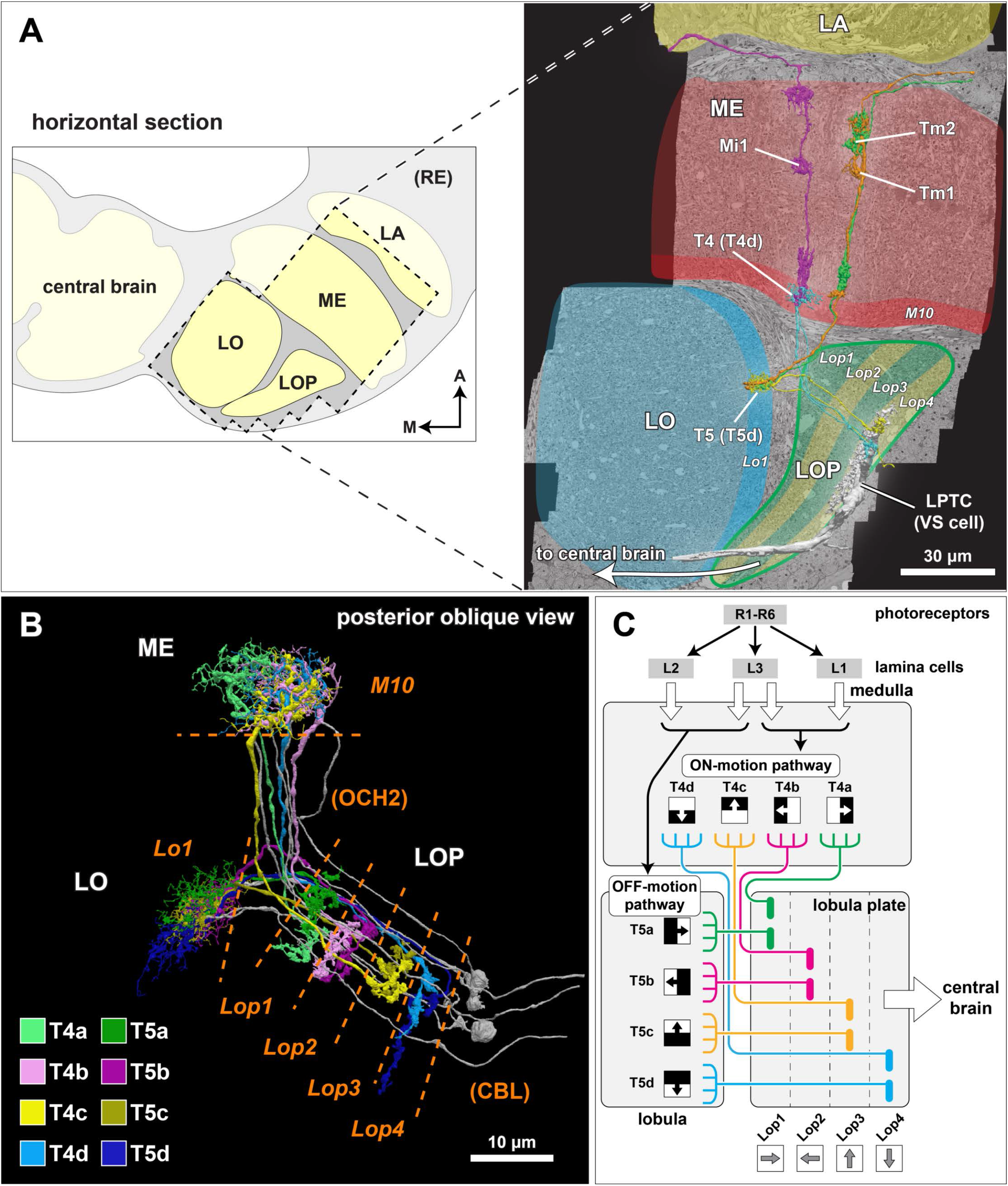
EM reconstruction of the synaptic partners of T4 and T5 cells in the lobula plate. (A) The optic lobe FIB-SEM dataset covers a subvolume of the medulla (ME), lobula (LO), and lobula plate (LOP), as well as the proximal part of the lamina (LA), selected to contain many connected neurons of the motion pathway. The data set was imaged with voxel size x = y = z = 8 nm, and the size of the image stack is 19,162 × 10,657 × 22,543 pixels, equivalent to 153 μm x 85 μm x 180 μm ^22^. In the right panel, representative neurons in the ON- and OFF-motion pathways in the medulla and the lobula, as well as a lobula plate tangential cell (VS cell) are shown (panel adapted from Shinomiya et al. ^22^). M: medial, A: anterior. (B) Subtypes of the T4 and T5 cells. The T4 cells receive inputs onto their dendrites in medulla layer 10 (M10), T5 neurons receive dendritic input in lobula layer 1 (Lo1). Both cell types project through the non-synaptic second optic chiasm (OCH2) and stratify into the four layers of the lobula plate (Lop1-Lop4). The cell bodies are located at the cell body layer (CBL) in the lobula plate cortex. The cell bodies and the cell body fibers are shown in gray, while some cell bodies are not shown. (C) A schematic diagram of the motion circuit. Local luminance is detected by the photoreceptors R1-R6 in the retina. The signals are relayed to the lamina cells (L1, L2, and L3), which send outputs to various columnar cells in the medulla (not detailed here). The 4^th^ order T4 and T5 neurons integrate inputs from the ON and OFF motion pathway neurons, respectively, and project to the lobula plate. The four subtypes (a, b, c, and d) detect visual motion in the front-to-back, back-to-front, upward, and downward directions, respectively, and project axons to the corresponding LOP layer where these directionally selective signals are integrated by lobula plate neurons.

Small volume EM reconstructions have revealed the synaptic connectivity of many neurons in the lamina, medulla, and lobula^18–23^, with special attention to the columnar neurons of the motion processing pathway. Together with functional studies, these reconstructions have revealed the detailed neuronal circuitry and the likely mechanism(s) of motion detection by T4 and T5 neurons. T4 are the ON directionally selective neurons. They encode the direction of motion, while none of their dendritic inputs (in medulla layer M10) do^24^. T5 are OFF directionally selective neurons that encode the direction of moving dark edges, by integrating inputs onto their dendrites in the first lobula layer (Lo1)^22,25^. Both cells have four distinct subtypes, a, b, c, and d. Each subtype projects axons to one of the four layers of the lobula plate (Figure 1B), where their terminals are retinotopically arranged^8,14^.

The lobula plate is the fourth neuropil in the optic lobe, and the evolutionary origin of this conserved neuropil has been hypothesized to relate to the origin of insect flight^26^. In Diptera (flies), the neuropil is best known for containing the dendrites of the ‘giant’ lobula plate tangential cells (LPTCs) that respond to specific patterns of visual motion^15,27–30^. The vertical system (VS) and horizontal system (HS) cells are the best studied LPTCs, and homologous neurons have been identified in both larger flies and *Drosophila*^14,28^. The arborization patterns of HS and VS cells in the *Drosophila* lobula plate were examined using the GAL4-UAS system and single cell labeling, confirming neuron morphology that closely resembles the corresponding cells of larger flies^31^, and the electrophysiologically measured response properties of these cells to visual motion patterns match those of larger flies^32,33^.

Based on imaging T4/T5 responses in the lobula plate^8^, it is now understood that each layer integrates inputs corresponding to one cardinal direction of motion: front-to-back (Lop1), back-to-front (Lop2), upward (Lop3), and downward (Lop4). Anatomical and physiological data suggest a correlation between an LPTC’s visual motion responses and its lobula plate layer pattern^28,34^. Further details of the lobula plate circuitry have not been thoroughly investigated, with the noteworthy exception of two bilayer lobula plate intrinsic (LPi) cells: LPi3-4 receive input in Lop3 and provide output to Lop4, while LPi4-3 sends signals from Lop4 to Lop3^14,35,36^. These LPi cells have been shown to inhibit their target LPTCs in response to ‘opponent’ motion. This sharpens the flow-field selectivity of the tangential cells, in a computation termed ‘motion opponency’^35^. Functional studies^3,35,37^ suggest that the site of action is the integration of excitatory, cholinergic T4 and T5 input together with inhibitory, glutamatergic LPi inputs by LPTCs, but the synaptic connectivity proposed by this parsimonious circuit hypothesis has not been verified.

The lobula plate also houses processes of columnar neuron types other than T4 and T5, including optic lobe-intrinsic neurons, such as Y, TmY (transmedulla Y), and Tlp (trans lobula plate) cells, which connect different optic lobe neuropils, and the LPC (lobula plate columnar), LLPC (lobula-lobula plate columnar), and LPLC (lobula plate-lobula columnar) cells, which are visual projection neurons (VPNs) into the central brain^3,14,38–44^. Detailed connectivity information for the principal neurons of the lobula plate, especially T4 and T5, LPTCs, LPis, and other columnar neurons, is largely unknown, and represents the last piece of the puzzle for the anatomical description of the primary motion information-processing circuit in the optic lobe. To close this gap, we reconstructed the neurons downstream of T4 and T5 in the lobula plate using an optic lobe dataset imaged with focused-ion beam-aided scanning electron microscopy (FIB-SEM)^22,45^. We exhaustively identified and cataloged T4 and T5 synaptic partners, and investigated complete synaptic profiles of the LPi cells that connect two layers of the lobula plate, as well as the HS and VS cells. In the process, we identified new cell types and attributed new connectivity patterns to known cell types, resolving several open questions about lobula plate connectivity, while also establishing many new neurons as important components of the motion pathway.

## Results

### EM reconstruction of the synaptic partners of T4 and T5 cells in the lobula plate

Our FIB-SEM data volume^22,45^ includes large parts of the lamina, medulla, lobula, and lobula plate (Figure 1A), covering regions corresponding to the eye’s equator, but not including the neuropils serving dorsal and ventral eye regions. Importantly this volume contains many connected neurons, corresponding to common retinotopic coordinates, enabling circuit reconstruction across these neuropils. Medulla neurons, including Mi1, Tm1, and Tm2, relay signals from lamina cells to T4, in M10, and T5, in Lo1 (Figure 1A)^22^. The four subtypes of T4 and T5 send outputs to one of the four LOP layers (defined to encompass the terminals of groups of T4 and T5 cells, see Methods), where they synapse with other optic lobe interneurons and VPNs leading to the central brain (Figure 1B,C).

### Connectivity of the seed T4 and T5 cells in the lobula plate

We reconstructed and then identified many neurons in the FIB-SEM volume, focusing on T4 and T5 cells and their targets. 277 T4s (66 T4a, 69 T4b, 74 T4c, 68 T4d) and 277 T5s (68 T5a, 74 T5b, 71 T5c, 60 T5d) were identified and at least partially reconstructed. Five cells of each subtype from a retinotopically overlapping region near the volume center were completely traced^22^. In the prior study, we detailed the dendritic inputs of these neurons, and here we describe the connectivity of these same 40 cells in the lobula plate. All computationally predicted synapses (see Methods) of these cells were proofread to identify their pre- and post-synaptic partners.

The connectivity of the inputs and outputs of the representative T4 and T5 cells in the lobula plate is summarized in Figure 2A, including all neurons connected with ≥5 synapses to any of the seed T4 or T5 cell (detailed connectivity data in File S1). We found 56 putative connected neuron types (mean of ≥5 synapses with any T4 or T5), including unidentified fragments (Figure 2A; shown in gray). 43 of these (77%) communicate with the same subtype of T4 and T5 within a single layer, resulting in a connectivity diagram that is largely comprised of four clusters, each corresponding to synapses within one lobula plate layer. One noteworthy exception is LPLC2, which is the only neuron we identified that receives inputs from all four T4 and all four T5 subtypes, corroborating the observations of a previous study that showed this cell type integrates spatially patterned inputs to selectively encode visual looming^3^.

**Figure 2.**
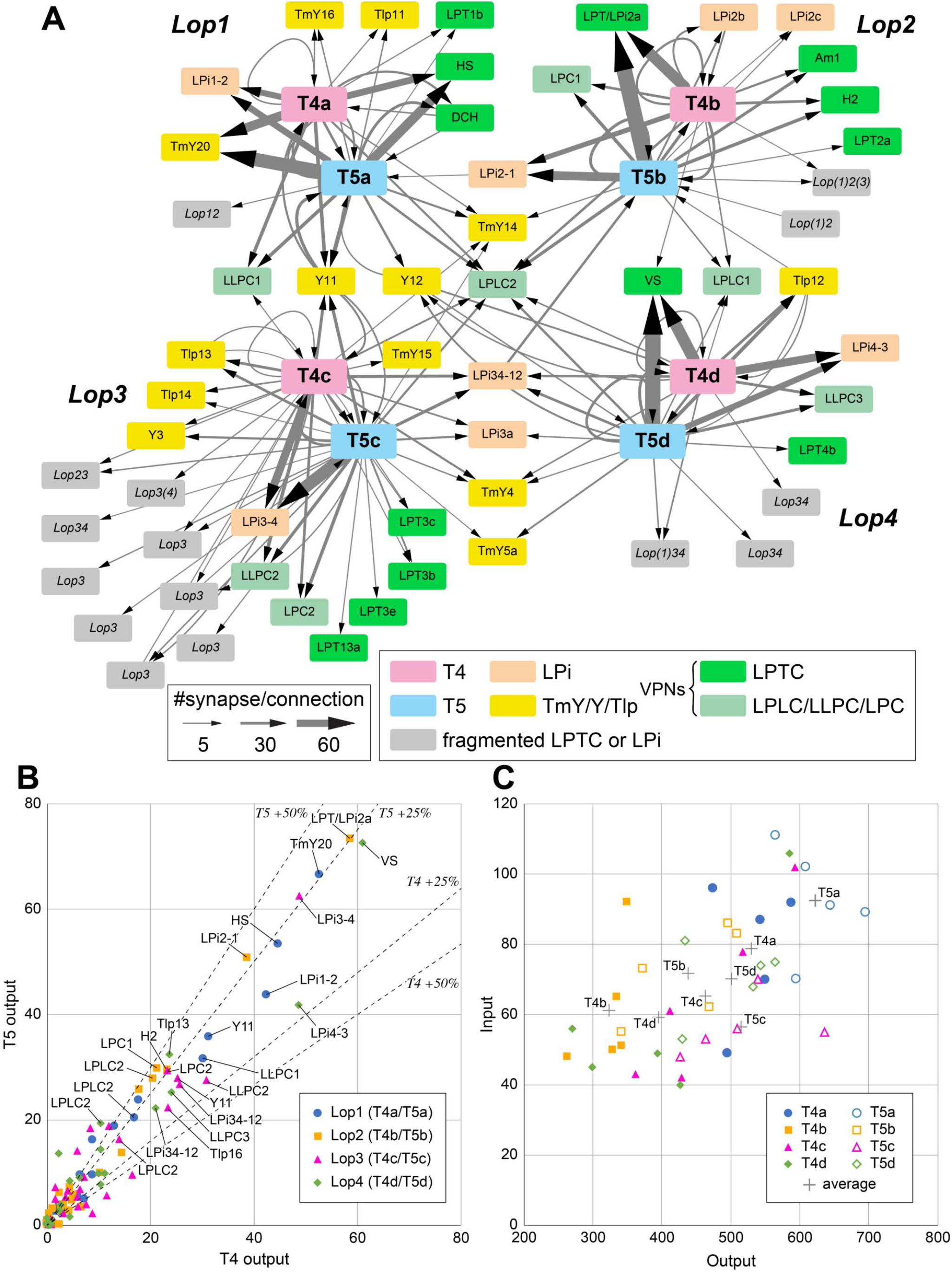
Connectivity of the seed T4 and T5 cells in the lobula plate. (A) The inputs and outputs of representative T4 and T5 cells (five cells per each subtype; see text for details, also File S1) in the lobula plate were comprehensively identified. The input and output cells were grouped by the cell type, and inputs and outputs corresponding to a mean of more than five synapses per T4 or T5 cell are shown in the diagram. The thickness of the arrows indicates the average number of synapses per T4 or T5 cell. Each rectangle indicates a cell type; colored rectangles correspond to uniquely identified cells, and gray rectangles represent neurons we could not uniquely identify due to incomplete reconstruction. For unidentified neurons, the main innervated layers are shown in italic letters. For example, *Lop(1)34* means that the fragment has major arbors in Lop3 and Lop4, and minor arbors in Lop1. LPi are lobula plate intrinsic cells, the TmY/Y/Tlp neurons connect the optic lobe neuropils, and LPTC and LPLC/LLPC/LPC cells are visual projection neurons (VPNs) that send outputs to the central brain. (B) Average numbers of output synapses from single T4 and T5 per postsynaptic cell type. Neurons are color-coded by the layer where they receive inputs from T4 and T5. Generally, outputs from T4 and T5 (and therefore inputs to their target neurons) are approximately evenly integrated by the postsynaptic cells, with a slight bias for T5. All named neurons receiving more than an average of 20 synapses from both T4 and T5 are labeled (LPLC2 is labeled for all four layers). The dashed lines indicate 25% and 50% difference from equal numbers of output from T4 and T5 to any target cell type. (C) Total numbers of input and output synapses of the representative T4 and T5 cells. Autapses (self-synapses) and synaptic contacts with glia are excluded from this quantification. Averaged synapse numbers of each cell type (five individual neurons per each cell type) are indicated as gray crosses.

How are the ON and OFF pathways integrated by targets in the lobula plate? In nearly every case, neurons that are strongly connected to T4/T5 have approximately symmetric inputs from T4 and T5, with a slight bias for T5 (pooled across all downstream neurons: 45.4% T4 vs. 54.6% T5), indicating that no major targets selectively integrate from only T4 or T5 (Figure 2B). This balanced integration of ON and OFF pathways suggests that any lobula plate neurons that primarily integrate inputs from T4 and T5 should not exhibit strongly asymmetric responses to bright vs. dark moving edges. However, some neurons may show differential sensitivity to dark or bright objects from other inputs. For example, LPLC2 responds strongly to dark looming stimuli and only weakly to bright looming^3^, despite substantial inputs from all T4/T5 subtypes (Figure 2B).

In mapping computational models onto the anatomy of the motion pathway, the T4/T5 axon terminals are treated as purely output structures^35^. We find that the axon terminals of T4 and T5 are primarily sites of synaptic output, but have some inputs: 87% of T4’s and 88% of T5’s lobula plate synapses are presynaptic (T4: 1765.2 pre/cell, 270.0 post/cell; T5: 2125.4 pre/cell, 295.4 post/cell). There are relatively small numbers of T4-T4, T5-T5, and T4-T5 connections within each layer. These occur between neighboring axon terminals, and each inter-terminal connection is typically ≤3 synapses. The number of pre- and postsynaptic sites per T4 and T5 varies by layer and individual neurons, but roughly follows a monotonic relationship; neurons with more output synapses tend to have more inputs (Figure 2C). For example, T4a and T5a neurons had more pre- and postsynapses than the other subtypes, due to strong connections with Lop1 neurons (Figure 2A, File S1).

T4 and T5 cells provide strong inputs to a diverse set of VPNs, of which many are large tangential cells (identified cells named and indicated in green; Figure 2A). We focus on the connectivity of the well-known HS and VS cells^28,31^ in Figure 3. Small-field VPN types (LPC, LLPC, and LPLC cells)^38,40,43,44^ are also found with substantial T4/T5 inputs in each layer. The morphology of many connected VPNs is shown in Figure 5. T4 and T5 cells in each layer provide strong inputs to four types of bilayer LPi cells, further explored in Figure 4. In addition to these cells, we identified many connections between T4/T5 neurons and other intrinsic optic lobe neurons, such as the TmY, Y, and Tlp cells (morphology shown in Figure 6) that interconnect different neuropils. For most newly described optic lobe intrinsic cell types, we provide light microscopy (LM) images as additional validation (Figures S1, S2). We summarize the core connectivity motifs at this output stage of the visual motion pathway in Figure 7.

**Figure 3.**
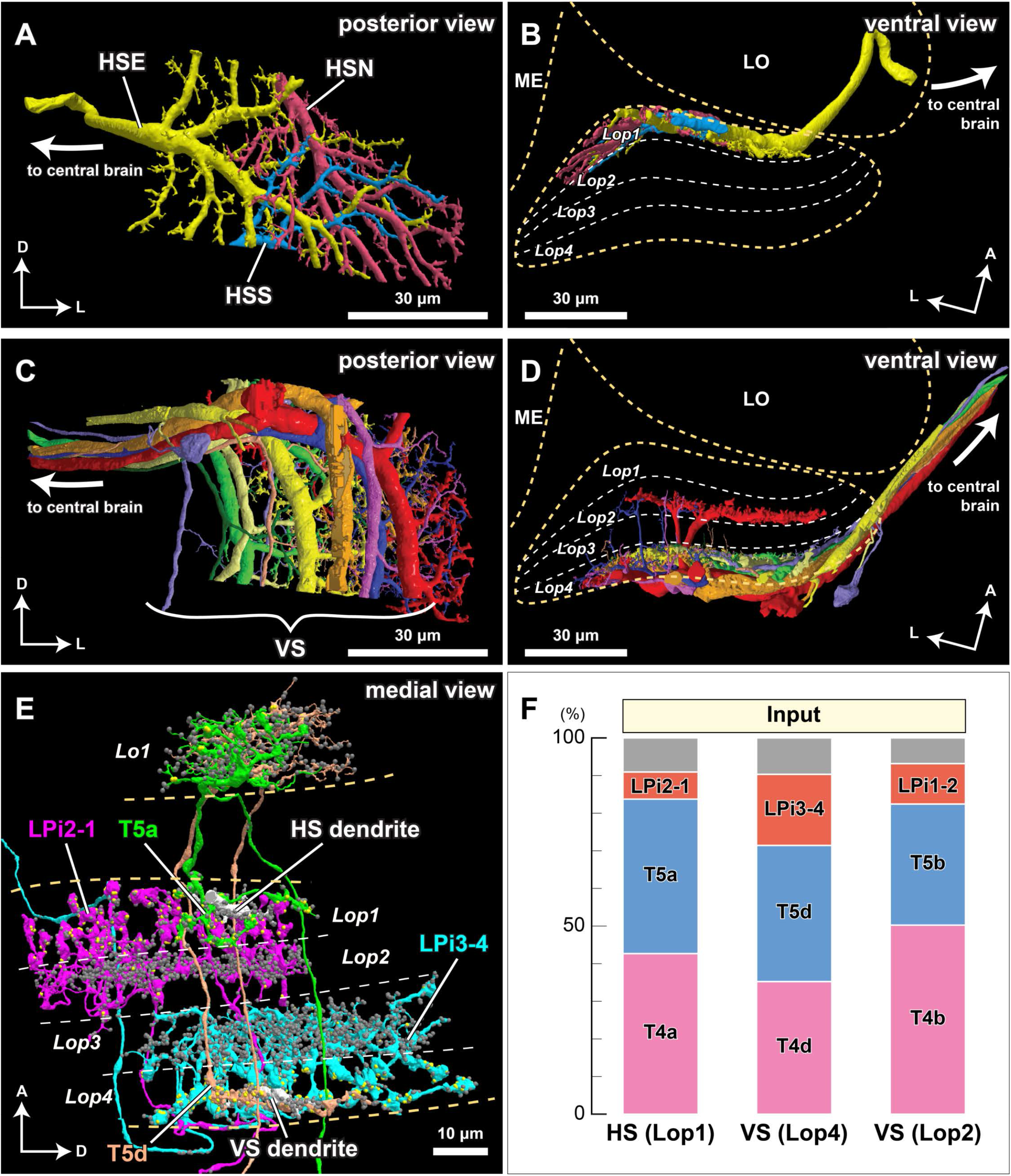
Synaptic connections of the horizontal system (HS) and vertical system (VS) lobula plate tangential cells. (A, B) The three HS cells (HSN, HSE, and HSS) occupy Lop1, the fist layer of the lobula plate. Collectively the dendrites of these neurons span Lop1 and overlap in the region of the lobula plate within our data volume, but are cut off at the edges of the volume. (A) posterior view, (B) ventral view. (C, D) The ten identified VS cells in our data volume. All have postsynaptic terminals in Lop4, while four of them also have branches in Lop2. (C) posterior view, (D) ventral view. (E) Examples of major input neurons to the HS and VS cells in the lobula plate. Single dendritic arbors (length ~20 μm) of one HS cell and one VS cell are shown in white. HS dendrites primarily receive input from the T4a, T5a, and LPi2-1 cells in Lop1, whereas VS dendrites in Lop4 primarily receive input from the T4d, T5d, and LPi3-4 cells. The T4 terminals are not shown to minimize clutter. Yellow and gray dots represent pre- and postsynaptic sites, respectively. (F) Inputs to the HS and VS cells. Synapses are verified and counted for small pieces of the HS and VS arbors in the respective layers (two branches for each of HS and VS (Lop4) and one branch for VS (Lop2)). Almost 90% of the inputs to the HS and VS cell dendrites come from T4, T5, and the bilayer LPi cells. A similar input distribution is found for the VS cells’ branches in the Lop2 layer, where they receive inputs from the T4b, T5b, and LPi1-2 cells. Gray indicates other, more weakly connected neurons or unidentified neuron fragments, less than 10% of the total synapses (detailed in File S2). No output synapses were found on these branches. The scale bars are approximate as the neurons are three-dimensionally reconstructed.

**Figure 4.**
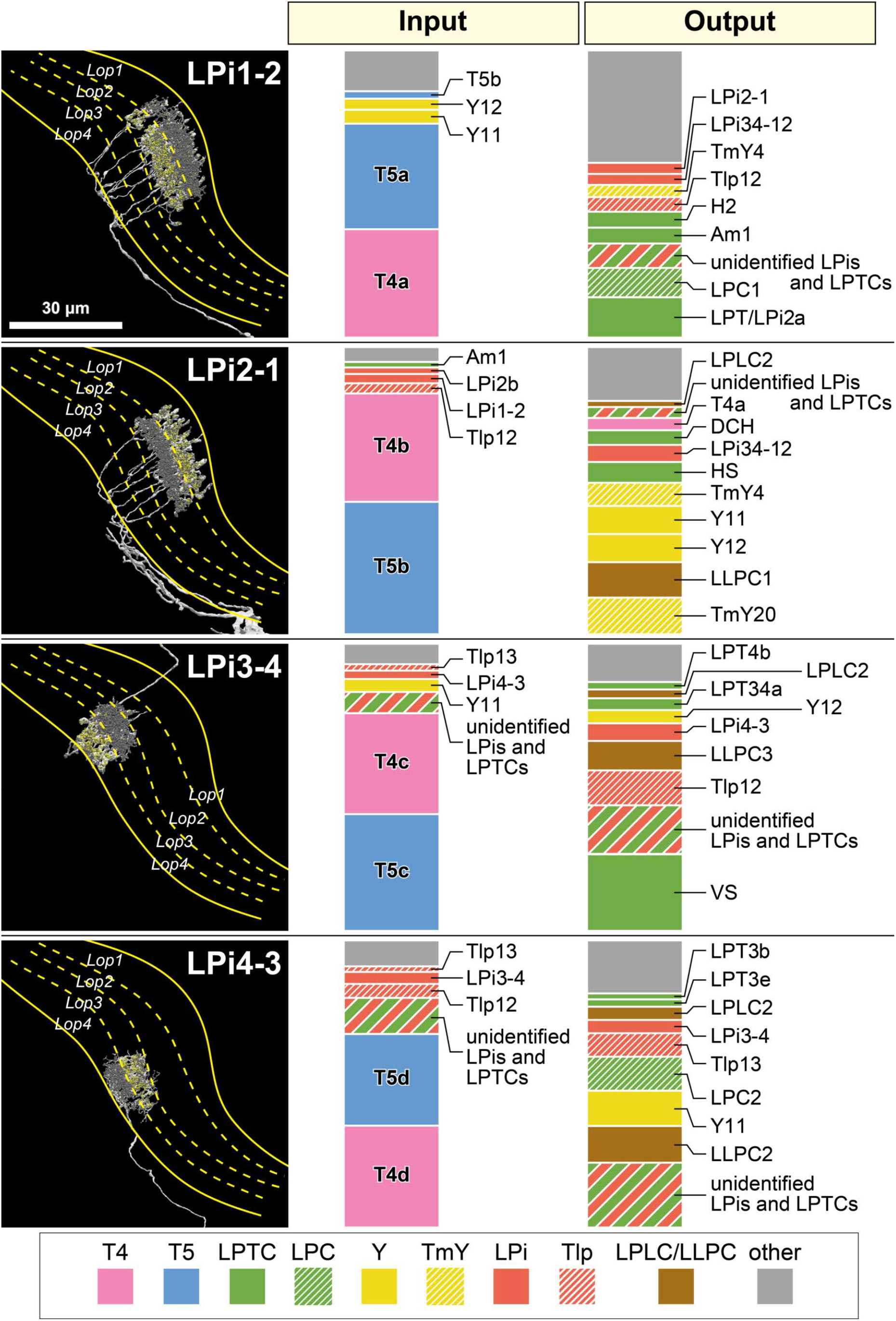
Connectivity of the bilayer Lobula Plate intrinsic (LPi) cells. A representative cell of each neuron type is shown in the left panel. Presynaptic sites are indicated with yellow dots and postsynaptic sites are shown with gray dots. These neurons primarily integrate inputs in one layer and supply outputs to the adjacent layer. Only the LPi3-4 cell is completely reconstructed, while the other cells are only partially reconstructed, since single neurons cover larger LOP areas than the imaged data volume. A candidate light microscopy match for LPi1-2 (Figure S1) suggests the possibility that the LPi1-2 reconstructions (and perhaps also the similar LPi1-2 fragments) may be parts of one or a few large cells. In the right two panels, ratios of the input and output synapses are shown for each indicated cell type. These data are based on a single selected branch for each cell type (with 600-1000 postsynaptic sites, 100-170 presynaptic sites), for which the pre- and postsynaptic connected neurons were identified wherever possible. Cell types occupying less than 2% of the total input or output synapses are not shown and are included as “other”. A number of tangential elements that have synapses with the LPi cells were only partially reconstructed due to the restricted data volume. These fragments of considerable size are grouped as “unidentified LPis and LPTCs”. Data summary based on File S3.

**Figure 5.**
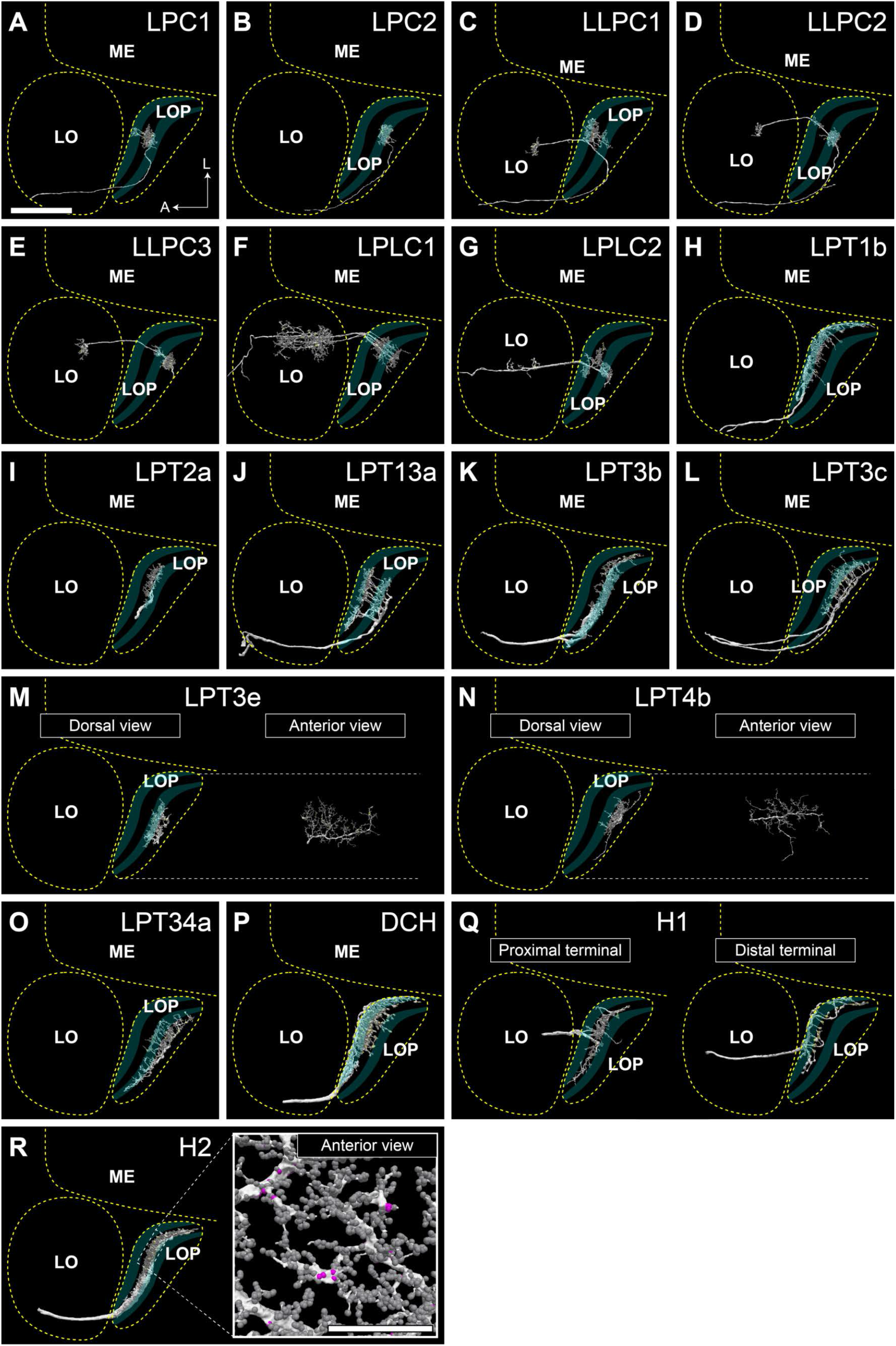
Lobula plate Visual Projection Neurons (VPNs) that integrate T4 and T5 inputs. The neurons are seen from the dorsal direction (horizontal projection), the approximate neuropil boundaries are outlined, and the LOP layers are indicated. Only neurons mentioned in other figures or in the main text are shown here. Some neurons are not fully reconstructed, especially the cell body fibers and the main axons projecting to the central brain. Lobula plate-lobula columnar (LPLC) cells have cell bodies in the cell body rind between the optic lobe and central brain and dendritic arbors in the lobula that extend into the lobula plate^38,40^, and project to the central brain from the lobula. Lobula plate columnar (LPC) and lobula-lobula plate columnar (LLPC) cells have cell bodies in the cell body rind of the lobula plate^38,40,41,43,44^, and project axons along a path posterior to the lobula plate to glomeruli in the posterior lateral protocerebrum. Both LPC and LLPC send a branch into the lobula plate which, in the case of LLPC cells, further extends into the lobula. HS and VS cells are omitted from this figure (see Figure 3). In (R), the branching pattern and synapse distribution is shown in the inset. Pre- and postsynapses are shown in magenta and gray, respectively. In (A), A: anterior, L: lateral. Scale bar: (A-S) 30μm, (R) inset 20μm.

**Figure 6.**
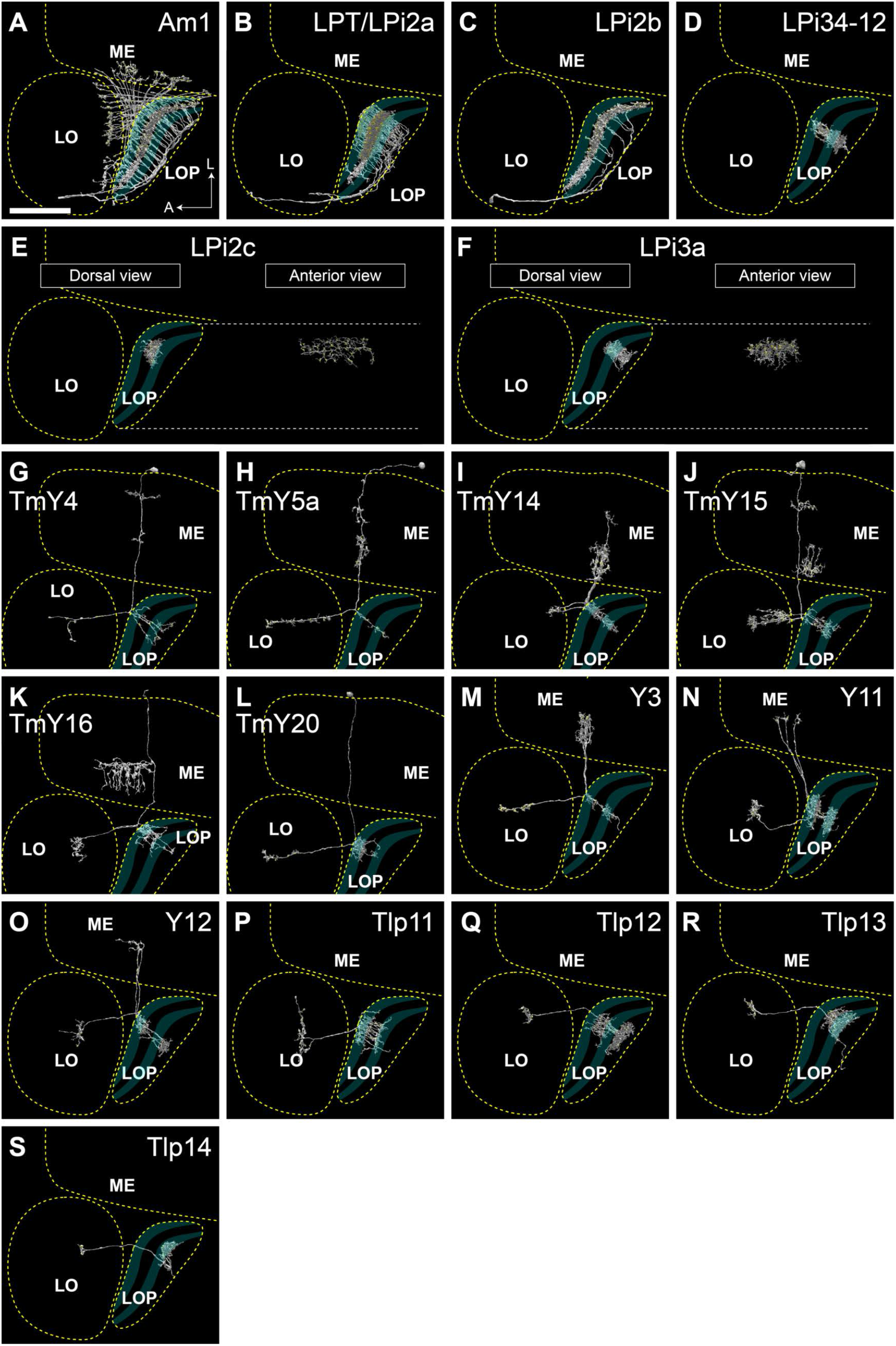
Optic lobe intrinsic neurons that integrate T4 and T5 inputs. The neurons are seen from the dorsal direction (horizontal projection), the approximate neuropil boundaries are outlined, and the LOP layers are indicated. Only neurons mentioned in other figures or in the main text are shown here. Some of the neurons are not fully reconstructed, especially the cell body fibers. We confirmed the general cell shapes of all newly identified cell types shown in this figure (with the exception of the comparatively small LPi2c and LPi3a fragments) by comparison to light microscopy images (Figures S1 and S2). Based on these matches, LPT/LPi2a may be a type of VPN with a central brain projection. Pre- and postsynapses are shown in magenta and gray, respectively. Scale bar: 30μm.

**Figure 7.**
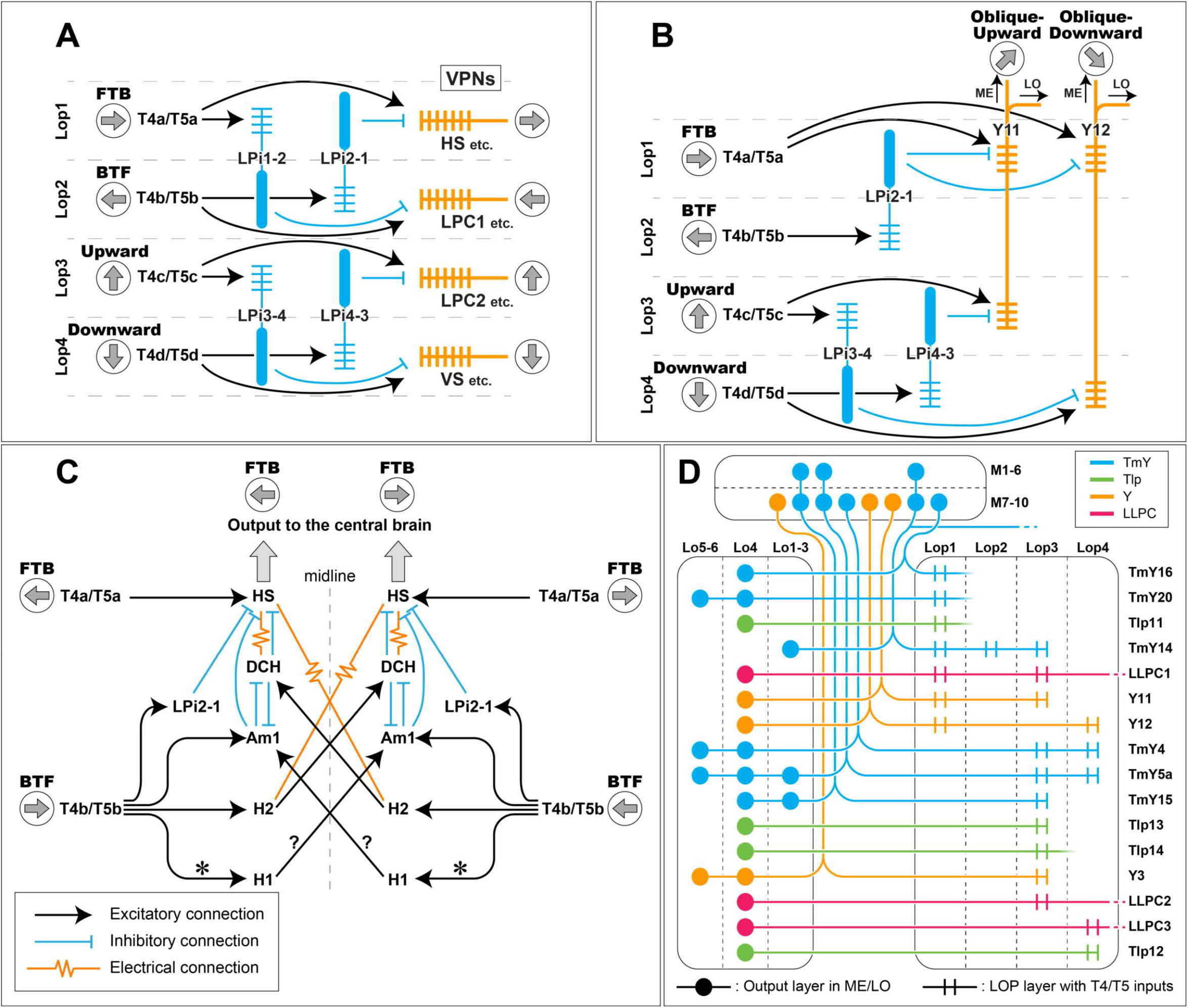
Summary of the motion pathway circuitry revealed by the lobula plate reconstruction. (A) The primary connections between the T4/T5 cells, bilayer LPi cells, and output VPNs. FTB: front-to-back, BTF: back-to-front. The LPi cells, indicated in blue, are likely all inhibitory cells (based on ^35^). Each layer has VPN outputs that are predicted (or known for the few supported by functional studies) to encode motion with the same directional selectivity as the T4/T5 subtypes in that layer. Some VPNs, like the VS cells of Figure 3, integrate inputs from multiple T4/T5 subtypes in different layers. (B) The Y11 and Y12 cells and their LOP inputs. The two Y cell types receive excitatory inputs from T4 and T5 in two layers and (putative) inhibitory inputs from T4 and T5 neurons in the other layers, via bilayer LPi cells. By integrating these inputs, it is expected that these neurons become most sensitive to the direction of overlapping sensitivity of their inputs, and thus Y11’s preferred direction would be for oblique-upward motion and Y12 would prefer oblique-downward motion. (C) The bilateral circuitry comprised of horizontal-motion sensitive neurons, including H1, H2, DCH, and Am1 cells, integrating motion from both eyes. Connections within the optic lobe are based on the observation of this dataset, while contralateral projections and synaptic contacts in the central brain are also based on prior studies and datasets, including ^43,59,61^. Connections from T4b/T5b to H1, indicated with asterisks (*), are not shown in Figure 2A, as the synapse numbers per representative were below the threshold, but shown in this diagram since the main inputs to H1 are T4b and T5b. H1 is considered to be a glutamatergic cell ^55^, and it is not known whether the signal from H1 is excitatory or inhibitory (see the main text for the detail). Since electrical synapses cannot be directly observed in the FIB-SEM dataset, the indicated connections are based on physical proximity of the axons of these cells in this dataset. The diagrams do not exhaustively list inputs and outputs of the shown neurons. (D) Neurons relaying T4/T5 outputs to the lobula. The paired parallel lines indicate T4/T5 inputs in the lobula plate, whereas the dots represent locations of the output (presynaptic) terminals in the lobula and the medulla. Input (postsynaptic) terminals in the lobula and the medulla, as well as terminals not coupled with T4/T5 in the lobula plate, are omitted. The central brain projections of TmY14 and the LLPC cells are shown as broken lines.

### Synaptic connections of the Horizontal System (HS) and Vertical System (VS) lobula plate tangential cells

The HS and VS cells are prominent LPTCs whose response properties have been extensively studied^31–33,46,47^. They represent major T4/T5 targets in their respective layers (Figure 2A). In Lop1, T4a and T5a provide strong inputs to the HS cells, with a mean of 44.6 synapses from each T4a and 53.4 synapses from each T5a (File S1). Each lobula plate houses 3 HS cells, HSN, HSE, and HSS (north, equatorial, and south) cells, which respectively cover the dorsal, middle, and ventral parts of the visual field^31^. In our imaged volume, we find identifiable fragments of all three cells (Figures 3A,B). The dendrites of these cells are almost purely postsynaptic. Based on the computational predictions, HSN, HSE, and HSS respectively had 6151, 4514, and 2066 postsynaptic densities (PSDs), and 7, 2, and 3 presynaptic T-bars. Most inputs to HS cells are from T4a and T5a (Figure 3F).

In Lop4, the VS cells receive a large portion of T4d and T5d’s synaptic outputs (Figure 2A,B). We identified 10 VS or VS-like cells in Lop4 (Figure 3C, D). This number exceeds the expected count of VS cells in *Drosophila* based on earlier genetic labeling studies^31^, but is consistent with the number in larger flies^30,48,49^ and a recent reconstruction in *Drosophila*^34^. These 10 cells have many common features: primary dendritic processes in Lop4, processes that are predominantly postsynaptic (typically >95% of total synapses), and simple connectivity profiles, with ~90% of inputs supplied by only three cell types (T4d, T5d, and LPi3-4; Figure 3F and Movie S1). Among the 10 VS and VS-like cells, four cells also have dendritic branches in Lop2 (Figure 3D; some cells could feature branching in other layers outside of the volume). VS cells with dendritic arbors outside Lop4 have been previously described^34,50^. Boergens and colleagues identified six VS and three VS-like cells in their dataset^34^, of which eight had branches outside of Lop4. Integrating directionally selective inputs in other layers is expected to shape the flow-field selectivity of these neurons to incorporate regional horizontal motion that accompanies body and head rotation around certain axes^47,50,51^.

The HS and VS cells are almost purely postsynaptic in the lobula plate. This connectivity from a very small set of cell types outlines a minimal number of circuit elements that could participate in the nonlinear summation of dendritic inputs by the HS and VS cells^52^. To quantify the input connectivity of these large neurons, we selected small (≤300 PSDs) branches of HS cells (one each from HSN and HSS) and VS cells (two Lop4 branches and one Lop2 branch; each from different cells) and proofread all synaptic sites. An HS branch and a Lop4 VS branch are shown in Figure 3E. A VS branch and its input neurons are shown in Movie S1. The summary of these connectivity analyses (Figure 3F) shows that ~80% of the input synapses of these cells are supplied by T4/T5. The HS (Lop1), VS (Lop4), and VS (Lop2) branches respectively receive 7.14%, 18.9%, and 10.7% of the input synapses coming from bilayer LPi cells (Figure 3F, File S2). Intriguingly, the Lop2 VS branch receives inputs mainly from T4b, T5b, and LPi1-2 cells, suggesting it indeed receives back-to-front local motion signals. Overall, the connectivity pattern between the T4/T5, LPis, and the giant LPTCs is very similar in these different layers (Figure 3E). This relatively simple connectivity structure strongly supports the expectations of previous functional studies of HS and VS cells—they appear to mainly integrate directionally selective inputs that are reinforced with motion opponent inputs from LPi neurons^35^.

### Connectivity of the bilayer Lobula Plate intrinsic (LPi) cells

We identified four bilayer LPi neuron types as major T4/T5 targets (Figure 2A). A previous study described LPi3-4 and LPi4-3, and based on the functional importance of these neurons for motion opponency, speculated about the existence of all four types^35^. In this study, we have reconstructed and identified LPi1-2 and LPi2-1 that bridge Lop1 and Lop2, confirming these prior predictions, although we are unable to describe the complete morphology of these neurons. We found a strong candidate for LPi1-2 using LM (Figure S1A). This apparent match suggests that LPi1-2, and perhaps also LPi2-1, may be considerably larger than LPi3-4 and LPi4-3. Confirming this proposal will require extensive reconstruction in a larger EM volume. All four LPi neuron types innervate two neighboring layers, with a stereotypic distribution of synapses (Figure 4, left). Each cell type has postsynaptic sites in one layer and presynaptic T-bars in the adjacent layer. At least 2/3 of the inputs are from layer-specific T4/T5 cells, while the outputs are shared by many neuron types (Figure 4, right; File S3). The LPi3-4 and LPi4-3 cells are glutamatergic^35,41^, and these cells provide inhibitory, directionally selective inputs to the target neurons^35,37^. Based on their similar morphology and connectivity, LPi1-2 and LPi2-1 are also likely inhibitory. This small circuit supports the proposed mechanism of Mauss et al.^35^: bilayer LPi cells integrate T4/T5 inputs in one layer and inhibit the postsynaptic neurons integrating the oppositely tuned T4/T5 signals in the adjacent layer, implementing motion opponency. The ~1/3 of LPi inputs provided by cells other than T4/T5 suggest that the lobula plate circuitry is more complicated, and perhaps more flexible, than the circuit models consider, but future connectomic and functional studies will be required to understand how these additional neurons contribute to motion processing.

### Lobula plate Visual Projection Neurons (VPNs) that integrate T4 and T5 inputs

In addition to the HS and VS cells (Figure 3), we identified several other VPNs as T4/T5 targets (Figures 2 and 5). In this study, we focused on identifying and quantifying T4/T5 target neuron connectivity, rather than describing complete synaptic profiles of the VPNs.

T4 and T5 connect with columnar VPNs, smaller cells that as a population cover large parts of the lobula plate. These cells belong to three main groups (LPLC, LPC, and LLPC) that are distinguished by cell body location, innervation pattern in the optic lobe, and axonal path to the central brain (further explained in Figure 5 legend). Based on their arbor sizes in the lobula plate and T4/T5 inputs, these cells are expected to respond to visual motion signals within small patches of the fly’s field of view. We distinguished two LPC types and three LLPC types based on lobula plate layer patterns (Figures 5A-E), in agreement with LM analyses^44^. LPC1 (Figure 5A), receives inputs from T4b and T5b. This anatomy suggests that these cells integrate back-to-front motion signals, which has been confirmed by calcium imaging^44^. LPC2 (Figure 5B) is a small-field VPN with T4c/T5c inputs (Figure 2A) and is therefore predicted to encode upwards motion. LLPC1 (Figure 5C), a VPN responsive to front-to-back visual motion^44^, has dendritic arbors in Lop1 and Lop3, with much stronger T4/T5 input in Lop1 (from T4a/T5a; Figure 2A). The synaptic terminal in the lobula appears to be mainly presynaptic (Figure 5C). LLPC2 and LLPC3 are similar cells with T4/T5 input in Lop3 and Lop4, respectively (Figures 5D-E). LPLC1 and LPLC2 cells^38,40^ are notable for receiving T4/T5 inputs in multiple lobula plate layers: T4/T5 a, b, c, and d for LPLC2, in agreement with the described mechanism of looming sensitivity in this cell type^3^ and T4/T5 b and d for LPLC1 (Figures 2A, 5F-G). By contrast, LPLC4^38,40^ is not a strong T4/T5 target in our dataset.

T4 and T5 neurons connect with LPTCs other than HS and VS cells, some of which we matched to known neurons, but in other cases, we name them based on their layer innervation patterns (Figures 5H-O). As many LPTCs are morphologically unique, we expect that many of these cells could be matched, one-for-one, to LM images or other EM reconstructions^34,53^. The dorsal centrifugal horizontal (DCH) cell (Figure 5P) is unique among this group as it is predominantly presynaptic to T4 and T5 (Figure 2A): 15.3% of T4a inputs and 12.7% of T5a inputs (excluding synapses between T4/T5 terminals) are from DCH, and it is by far the largest input to T4/T5 from a single LPTC. The terminals of DCH cover the dorsal half of Lop1, while the homologous ventral centrifugal horizontal (VCH) cell covers the ventral half^34,54,55^. The CH neurons innervate the ipsilateral inferior posterior slope (IPS) in the central brain, are GABAergic^55–57^, and likely inhibitory. Although we did not find VCH (due to the imaged area restriction), our data suggest that these two cells are the only major LPTCs that feed signals from the central brain to T4a/T5a (File S1).

H1 is a heterolateral LPTC directly connecting both lobula plates (Figure 5Q)^28^, and is sensitive to ipsilateral back-to-front visual motion, similar to H2^58,59^. We found two profiles that likely correspond to proximal and distal terminals of both H1 cells. The proximal terminal is predominantly postsynaptic and confined within Lop2, while the putative distal terminal branch is presynapse-rich, with boutons mainly in Lop1 and Lop2. T4b and T5b provide synaptic inputs to the proximal terminal, but H1 does not appear in Figure 2A since the averaged numbers of synapses per terminal (~4.8 from both T4b and T5b) were below our threshold for inclusion. Nonetheless, H1 is expected to integrate many inputs from T4b/T5b throughout Lop2, which is consistent with the described motion preference^28,54,58,59^. The distal terminal of H1 has limited synaptic contacts with T4 or T5 cells (only accounting for ~0.1% of H1’s predicted output synapses).

The H2 cell, another identifiable LPTC that is well-known from work in larger flies, has dense neuronal processes confined to Lop2 (Figure 5R) and projects to the IPS in the contralateral hemisphere of the brain^30,55^. Unlike HS or VS cells, H2 branches in Lop2 feature mixed pre- and postsynaptic terminals (Figure 5R, inset), as suggested by genetically-driven synaptic markers^55^. H2 reportedly connects with the CH cells in the central brain^58^, and a central brain EM connectome dataset (“hemibrain”) revealed that H2 provides the strongest input to DCH and VCH^43^. H2 is thus strongly coupled with the CH cells from the opposite brain hemisphere, contributing to processing motion information from both eyes.

### Optic lobe intrinsic neurons that integrate T4 and T5 inputs

T4 and T5 target optic lobe intrinsic cells in addition to the bilayer LPi neurons, including several types of LPi, TmY, Y, and Tlp neurons (Figure 6). We identified both known and new optic lobe intrinsic cell types as we described T4/T5 targets. For most of the new cell types in this group, we further confirmed the morphology with LM matches (Figure S2).

One noteworthy target is Am1 (Figure 6A), a single, large, amacrine-like neuron innervating the medulla, lobula, and lobula plate with tree-like arborization^27,45^. Am1 receives inputs from T4b/T5b in Lop2 (Figure 2A) and has significant synaptic contacts with some LPTCs. The predicted synapses contain strong inputs from DCH and contralateral H1, and outputs to DCH and HS cells. Based on these connections, we expect that Am1 is inhibitory (since it is unlikely to excite HS cells in response to ipsilateral T4b/T5b input) and participates in a bilateral circuit comprised of several tangential cells that integrate horizontal motion signals from both eyes^58–61^ (Figure 7C).

We find several putative LPi and LPi-like neuron types (Figures 6B-F) that all differ from the bilayer LPi types and provide further examples of the diverse neuronal composition of each layer. A large cell we tentatively named LPT/LPi2a receives the strongest inputs from T4b and T5b among all the neurons in our data (Figures 2A and 6B). LPT/LPi2a has a similar but distinct morphology from the bilayer LPi2-1 cell in the lobula plate, with main branches containing pre- and postsynapses in Lop2 with additional sparser processes in Lop1. While T4/T5 supply >80% of LPi2-1’s input, this number is <50% for LPT/LPi2a, suggesting it participates in circuits with more elaborate connectivity than the main bilayer LPis (Figure 7A). Our best candidate for an LM match is a VPN with a projection to the central brain (Figure S1B). LPi2b is another large Lop2 cell that appears to span the entire lobula plate, but with a more restricted layer pattern and fewer inputs from T4/T5 (Figures 6C and S1C). LPi34-12 (Figure 6C; named for its layer pattern) is similar to the bilayer LPi cells but receives T4/T5 input in both Lop3 and Lop4 and has output synapses in Lop1 and Lop2 (Figure 2A), and thus appears to represent an undescribed interaction between motion detected along different directions.

TmY cells have cell bodies in the medulla cell body rind and terminals in both the lobula and lobula plate (Figures 6G-L). TmY4, TmY5a, TmY14, and TmY15 have been previously described^14,20–22^, while TmY16 and TmY20 are reported here for the first time and confirmed with LM matches (Figures S2A-B). TmY20 has the highest number of inputs from T4a/T5a of all the targets we found (52.6 synapses/T4a and 66.6 synapses/ T5a; Figure 2A, File S1). Unlike most TmY cells, we don’t find synapses on the TmY20 neurite in the medulla; the cell synapses only in the lobula and lobula plate (reminiscent of LPi3-4, which also lacks synapses in the medulla^35^). TmY20 has mostly presynaptic terminals in lobula layers Lo5 and Lo6 (Figure 6L), suggesting this neuron relays front-to-back motion information to lobula neurons. The other TmY cells have extensive arborizations outside the lobula plate and a full inventory of their connectivity may be required for detailed predictions about their role in motion processing. Y cells (Figures 6M-O) are columnar neurons with cell bodies in the rind posterior to the lobula plate, that innervate the medulla, lobula, and lobula plate^14^. Tlp cells (Figures 6J-S, S2E-H) are similar to Y-cells but lack a medulla branch. We identify one known (Y3) and two previously undescribed Y neurons (Y11, Y12) as T4/T5 targets, and confirm their morphology using LM (Figures S2C-D). Tlp, Y and TmY cells all provide paths for relaying different subsets of retinotopic T4/T5 outputs to the lobula (Figure 7D).

The two Y-cell types identified here, Y11 and Y12, are notable for integrating T4/T5 input from different layers: Y11 from Lop1 and Lop3 and Y12 from Lop1 and Lop4. The two cells are otherwise morphologically very similar, with boutons in the same medulla and lobula layers. Both cell types have pre- and postsynaptic contacts with T4 and T5 (File S1), integrating their signals in their respective layers. Since Y11 synthesizes front-to-back (Lop1) and upward (Lop3) motion signals and Y12 combines front-to-back and downward (Lop4) motion signals, the two cells are likely to each encode a preferred motion direction along the oblique directions in-between the preferred cardinal directions of their input T4s and T5s (Figure 7B).

## Discussion

The giant tangential cells of the fly lobula plate have received considerable interest for decades^30,62,63^, but the descriptions of the circuits at this ‘final’ optic lobe stage of the motion pathway have been rather incomplete. In this study, we used 3D-EM reconstructions to inventory the synaptic partners of T4 and T5 neurons with completeness unmatched by other approaches. Our work reveals a much more elaborate architecture for processing visual motion, with several major new findings: 1) lobula plate target neurons integrate T4 and T5 inputs with approximately equal weights, 2) each layer houses a unique ensemble of downstream neurons, while sharing a core circuit motif composed of T4/T5, a bilayer LPi cell, and output VPNs, 3) new circuit elements that combine motion signals for different directions, including the Y11 and Y12 cells, and, 4) many neurons conveying motion signals from the lobula plate to the lobula, implicating lobula circuitry with a more significant role in motion processing.

We found that all lobula plate neurons that are strongly connected to T4 and T5 axon terminals integrate these inputs, in the same layers, with nearly equal weight (Figure 2B). This is a conceptually significant finding, as it implies, at least for the motion pathway, that the ON and OFF separation is an internal feature of the optic lobe, and at the output stages of the pathway, the ON and OFF motion signals are combined onto all prominent lobula plate targets.

Most of the identified LPTCs receive T4/T5 inputs in single layers, while three columnar VPNs (LLPC1, LPLC1, and LPLC2) and some optic lobe intrinsic neurons (e.g., LPi34-12) receive T4/T5 inputs in multiple layers (Figure 2A, 5, 6). These connectivity patterns suggest that most LPTCs carry large-field motion information representing one of the four cardinal directions, while small-field neurons may integrate signals from multiple layers and as a population could transmit more complex motion information to their downstream neurons. The best explored example of this is LPLC2, whose looming sensitivity was attributed to T4/T5 and bilayer LPi inputs in all four layers ^2,3^, a hypothesis that this study has substantively confirmed.

### Bilayer LPi cells

Most neurons we describe, such as the LPTCs or the columnar cells, appeared variable across the layers, only T4, T5, and the bilayer LPi cells exist in nearly identical, layer-specific subtypes. The four bilayer LPi cells have a common distribution of synapses, with nearly equal T4/T5 inputs in one layer, and substantial output synapses in a neighboring layer, where they presumably inhibit most or all of the neurons that also receive excitatory T4/T5 inputs in that layer (Figures 2A, 4, and 7A), implementing motion opponency^35^. While the bilayer LPi cell types likely serve similar functions in motion processing, there are also clear anatomical differences. The cell bodies of LPi1-2, LPi2-1, and LPi4-3 are in the lobula plate cortex, while LPi3-4’s are in the medulla cortex^14,22,35^, and therefore likely derive from different precursor cells. EM and LM data suggest that there are substantial size differences among the bilayer LPis, with LPi3-4 likely the smallest arborization, and individual LPi1-2 cells potentially arborizing across much of the lobula plate (Figure S1A). Since the spatial coverage of individual LPi neurons differs between the four types, the spatial integration of opponent signals may differ between layers, for reasons that are unclear and merit further investigation. These differences raise questions about the evolution of the bilayer LPi cells. Are these LPis derived from a shared ancestral cell type, for example via duplication, that later substantially diverged in some of their anatomical properties? Or did the antiparallel inhibition mediated by the bilayer LPis evolve independently in different layers?

### Y cells encoding oblique motion directions

We discovered that Y11 and Y12 integrate motion information in two layers and thus likely synthesize a preferred tuning for a new, oblique direction of motion (Figures 2A and 7B). These neurons effectively fill two gaps between the four cardinal directions represented by T4/T5 subtypes. Both neurons combine a vertical motion signal with front-to-back motion, but we did not find complementary neurons for the oblique motion directions integrating Lop2/back-to-front motion. This asymmetry may reflect a bias for motion components experienced during forward locomotion. The Y cells have some similarities but also large differences with LPLC2, as they receive T4/T5 inputs from spatially overlapping areas in different layers, and their main targets are in the lobula and medulla (Figures 6N,O, and 7D). Taken together, this suggests that Y11 and Y12 likely synthesize ‘new’ preferred directions of motion sensitivity which is then further processed or integrated with other visual modalities. Identifying the targets of Y11 and Y12 will be an important goal of future connectomes.

### Expanding the horizontal motion detection circuit with new cell types

Our detailed analysis of the neurons connected to T4/T5 in Lop1 and Lop2 suggests several new connections should be added to existing models of binocular integration of rotational optic flow derived from work in blowflies^61^. The Am1 cell, which receives inputs from ipsilateral T4b/T5b and contralateral H1, likely combines optic flow across both eyes. H1 expresses a marker for glutamatergic neurons^55^. In *Drosophila*, glutamate could function as either an excitatory or inhibitory transmitter, while in blowfly, H1 seems to provide excitatory signals^58^. T4b and T5b detect back-to-front movement, and via (putative) inhibitory LPi2-1cells, suppress the activity of neurons in Lop1, including HS cells (Figure 7C). Am1 may represent two more pathways for suppressing the activity of HS cells in response to back-to-front motion inputs, directly, and through DCH, which is also electrically coupled with HS in *Calliphora*^61^. It would appear that opponency is accomplished at different scales—the scale of bilayer LPi neurons and the CH neurons, and over the entire field of view by combining contralateral optic flow transmitted by H1 and H2 (Figure 7C).

### Multiple neuron types convey T4/T5 signals to specific lobula layers

Our analysis shows that T4/T5 have strong synaptic contacts with a variety of neuron types that appear to relay these signals within the optic lobe. For example, TmY20 cells (Figure 6L), receive the largest share of T4a/T5a output synapses (Figure 2A). While the standard circuit models of the motion pathways, comprised of T4/T5, LPTCs, and bilayer interneurons (Figure 7A), have remained compact, evidence for additional, strong pathways suggests a broader role for motion signals. A substantial fraction of T4/T5 downstream cells, including Tlp, LLPC, and TmY neurons (Figures 2, 5, and 6) project to the lobula, where they mainly target layer Lo4 (Figure 7D). The circuits of the lobula, outside of the T5 inputs in Lo1, have been scarcely examined. What is now clear is that motion signals passed from the lobula plate should significantly contribute to visual pathways in the lobula, and potentially many VPNs projecting to the central brain could inherit motion signals from the lobula plate without any input sites there. The complete description of these pathways and their extended circuits will require an EM data set that covers all neuropils of the optic lobe as well as the central brain.

### Towards complete reconstruction of sensory-to-motor pathways

The connectivity profile of T4/T5 in the lobula plate we present here fills a large missing part of the motion pathways, the link between the detection of directionally selective motion and visual projection neurons of the lobula plate. With this part finally reconstructed, the motion pathway from the photoreceptor cells to the central brain can now be traced neuron-by-neuron by combining the accomplishments of multiple 3D-EM reconstructions^18–20,22,23,64^. Many of the VPNs we reconstructed here are also identified in the hemibrain dataset^43^ that contains much of the central brain, enabling the comprehensive identification of downstream circuits to extend the described pathways even further. Many of the new discoveries reported here suggest a more integrative picture of optic lobe processing, where the lobula plate is no longer seen as the sole substrate for motion processing, but rather is understood to organize ON and OFF directionally selective signals for a variety of as-yet unexplored roles in visually guided behaviors.

## Acknowledgements

The authors thank members of the FlyEM Project Team at Janelia Research Campus for sample preparation, image acquisition, image processing, and proofreading of the neurons and synapses and the FlyLight Project Team for light microscopy images. We especially thank Stuart Berg, Lowell Umayam, and William Katz of the FlyEM Team for data management and preparing neuPrint/neuroglancer data, and Gerry Rubin and the members of the FlyEM steering committee for supporting this project. We also thank members of the Reiser lab for fruitful discussions and advice on the analysis. This project was supported by HHMI.

## Methods

### The EM dataset

All of the results presented in this manuscript were based on the same optic lobe FIB-SEM data volume that was used in two previous studies ^22,45^. The sample was obtained from the right optic lobe of a 6-day post-eclosion female fruit fly, *Drosophila melanogaster*, a cross between homozygous *w^1118^* and CS wild type. The tissue was imaged with FIB-SEM with an isotropic voxel resolution (x = y = z = 8 nm). The size of the image stack is 19,162 × 10,657 × 22,543 pixels, equivalent to 153 μm x 85 μm x 180 μm of the brain. The grayscale data of the image volume as well as the reconstructed neurons is available at http://emdata.janelia.org/optic-lobe/. Connectivity data will be made available through neuPrint, an online tool for accessing and analyzing connectome data ^65^. For more information, see the **EM reconstruction of synaptic partners of T4 and T5 cells in the lobula plate** section and our previous publication ^22^.

### Reconstruction of the neurons and the neuron nomenclature

Neuronal profiles were automatically segmented, and synaptic motifs (presynaptic T-bars and postsynaptic densities) were predicted throughout the volume as described previously ^22^. Predicted synapses reliably reveal connectivity of most neurons and polarity of most synaptic connections ^22^, while they include some false-positive and false-negative synapses. For the main connectivity results analyzed and presented here, we manually proofread all predicted pre- and postsynapses of the 40 core T4 and T5 neurons as well as the dendrite fragments of the HS and VS cells (Figure 3) and the bilayer LPi cells (Figure 4) for higher quality results. Neurons and synapses were proofread and visualized using the NeuTu ^66^ software package.

After identifying representative T4 and T5 cells, five cells per each subtype, their synaptic partners in the lobula plate were exhaustively traced, though not necessarily to completion. Most of the cells documented in previous studies, including prominent LPTCs, were identified by their morphology. When two or more neurons have similar morphology, information of the spatial distributions of pre- and postsynaptic terminals, synapse counts, as well as the neuron types sharing synaptic connections were used to determine the cell types. New neuron types identified in this work (part of Figures 5 and 6) were named following the nomenclature convention of the optic lobe neurons primarily introduced by Fischbach and Dittrich ^14^. The lobula plate tangential cells (LPTCs) have traditionally been given unique names, such as the HS, VS, and CH cells. Newly found LPTCs were distinguished by the extent of branching arbors in the lobula plate. Using a similar format used by Fischbach and Dittrich ^14^ and Otsuna and Ito ^67^ for other neuron types, we tentatively named these cells by combining LPT (lobula plate tangential) + innervating layers + alphabetical identifier, e.g., LPT3b and LPT34a. This nomenclature aligns with neuron names such as the lobula tangential (LT), medulla tangential (MT), and lobula columnar (LC) cells, while using “C” for “cell” was avoided for naming individual neurons as it is commonly used to abbreviate “columnar”. Likewise, the names for the columnar lobula plate cells, LPC, LLPC, and LPLC, match the names used in other studies carried out at the Janelia Research Campus ^38,43,44^. Neurons were given tentative names as far as the overall morphology was reconstructed or, at least, a characteristic branch in the lobula plate was sufficiently reconstructed (in the case of LPi and LPTCs). Numbers used in the names of the Tlp and Y cells were selected to avoid overlap with numbers in Fischbach and Dittrich ^14^ (since EM/Golgi matches can be inclusive). Gaps in the numbering of TmY neuron types reflect cell types identified in ongoing work that are not T4 or T5 synaptic partners by the criteria used in this study and therefore are not included here. In contrast to the bilayer LPi names, the names of the TmY, Tlp, Y cells, etc. do not refer to the lobula plate layer pattern of these neurons.

### Light microscopy (LM) and LM/EM comparison

Individual cells were labeled using MultiColorFlpOut (MCFO) ^16^. Details of the fly crosses for each supporting figure panel are listed in Table S1. All images show cells from female flies. Images were acquired on Zeiss LSM 710 or 780 confocal microscopes with 63 × 1.4 NA objectives at 0.19 μm x 0.19 μm x 0.38 μm or 0.38 μm x 0.38 μm x 0.38 μm voxel size. Samples were prepared and imaged by the Janelia FlyLight Project Team. Detailed protocols are available online (https://www.janelia.org/project-team/flylight/protocols). We used GAL4 lines from the Janelia and Vienna Tiles collections ^11,17^. Figures show views of substacks rendered using VVD viewer (https://github.com/takashi310/VVD_Viewer). In some cases, additional labeled cells or background signal were removed by manual editing in VVD viewer. Original confocal stacks will be made available online.

LM and EM matches are based on visually comparing anatomical features, in particular cell body location and arborizations in specific optic lobe subregions and layers. With the exception of LPi2c and LPi3a (which we did not attempt to match due to their comparatively few distinct features and small size) and LPi2-1 (for which we did not identify LM images), we confirmed the cell shapes of all newly identified optic lobe intrinsic cell types by identifying probable light microscopy matches.

---

**Figure S1.**
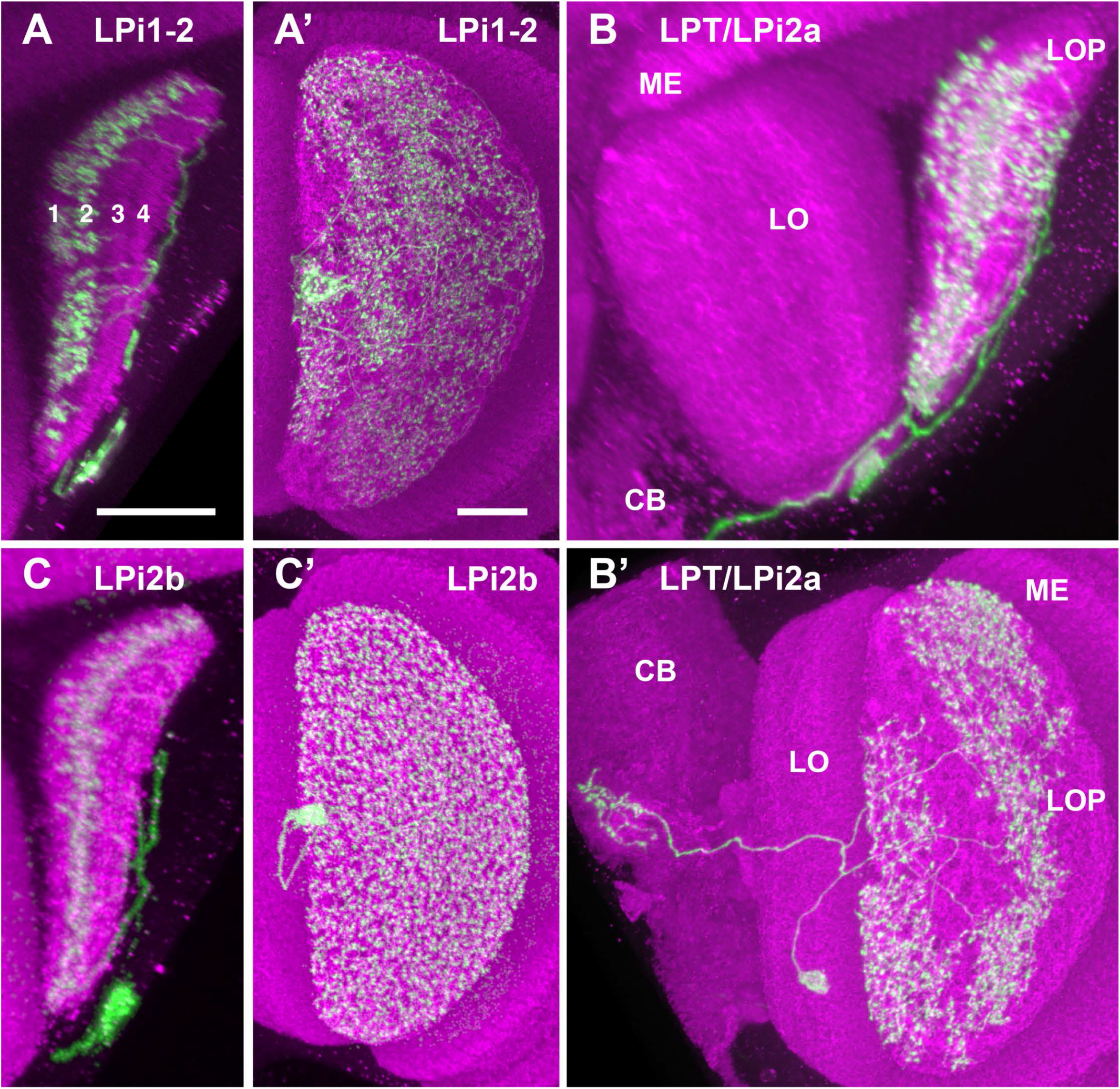
Candidate light microscopy matches for large LPi-like cells. Related to Figure 6. Images show resampled views generated from confocal stacks with MCFO-labeled neurons using VVD viewer (see Methods). Some images were manually edited in VVD viewer in order to only show the cells of interest. Panels show either the lobula plate layers (in a view similar to Figures 5 and 6) (A, B, C) or an *en face* view of the lobula plate from posterior (A’, B’ C’). Cell type names indicate apparent EM matches. Scale bars in A and A’ represent 20 μm. Other panels are shown at similar but not identical scale. Note that the neuropils appear to be much smaller than those in the EM sample (e.g., Figure 1A) due to shrinkage from dehydration for DPX mounting (see Methods and also Figure 19 of Scheffer et al. ^43^). Numbers in panel A mark lobula plate layers. Brains regions are indicated in (B, B’) (CB, central brain). (A) A large lobula plate intrinsic cell that locally matches the arbor structure (thin processes, likely dendritic in LOP1, varicosities, likely presynaptic, in LOP2; parallel processes to and soma in the lobula plate cell body rind) of LPi1-2 (Figure 4). A’) Arbor spread of the neuron in (A). Processes cover most of the LOP in a non-uniform pattern. (B) Layer pattern and lobula plate coverage (B’) of a neuron resembling LPT/LPi2a (Figure 6B). Central projection suggests that the neuron we identify in the EM volume as LPT/LPi2a is likely a VPN. (C) Layer pattern of an apparent LM match of LPi2b (Figure 6C). This cell covers all of the lobula plate (C’).

**Figure S2.**
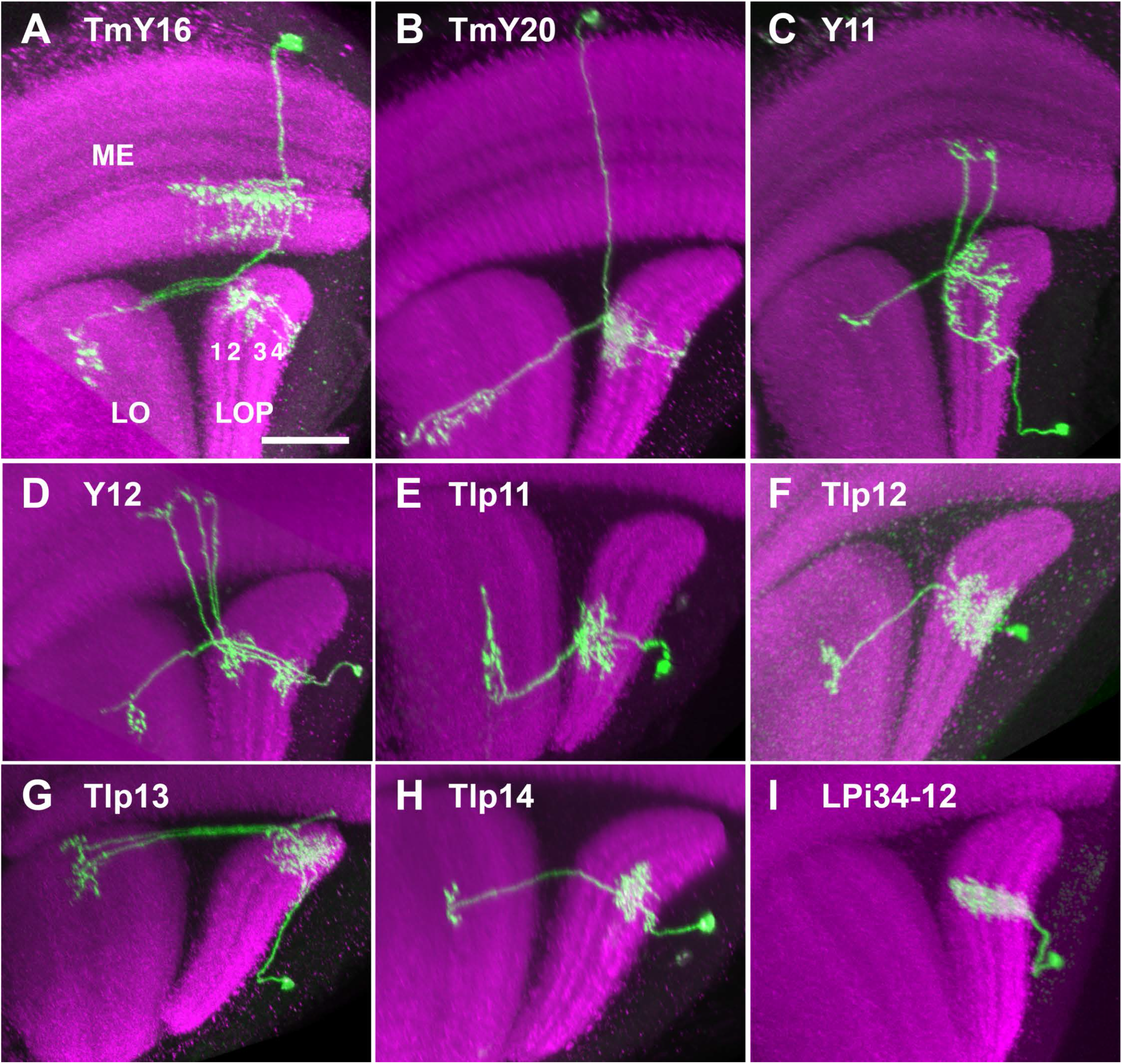
Candidate light microscopy matches for newly described optic lobe intrinsic cell types. Related to Figure 6. Images show resampled views generated from confocal stacks with MCFO-labeled neurons using VVD viewer (see Methods). Some images were manually edited in VVD viewer in order to only show the cells of interest. Panels show the lobula plate layers (displayed in a view similar to Figures 5 and 6). Cell type names indicate apparent EM matches. Scale bar in A represent 20 μm. Other panels are shown at similar but not identical scale. Numbers in panel A mark lobula plate layers, optic lobe regions as indicated. (A) TmY16, (B) TmY20, (C) Y11, (D) Y12, (E) Tlp11, (F) Tlp12, (G) Tlp13, (H) Tlp14, (I) LPi34-12.

## Supplemental Information

**Table S1:** Fly crosses used to visualize lobula plate neurons in this study.

| Figure panel(s) | Fly cross |
| --- | --- |
| S1A, S1A' | R20F11-GAL4 crossed to MCFO-7 (Nern et al 2015) |
| S1B, S1B' | R15D05-Gal4 crossed to MCFO-7 |
| S1C, S1C' | R42B05-GAL4 crossed to MCFO-7 |
| S2A, S2C, S2D, S2I | OL-KD (29C07-KDGeneswitch-4) in attP40; R57C10-GAL4 in attP2 tubP-KDRT>GAL80- 6-KDRT> in VK00027 crossed to MCFO-1 (Nern et al 2015) |
| S2B | VT048842-GAL4 crossed to MCFO-7 |
| S2E | R10E08-GAL4 crossed to MCFO-7 |
| S2F | R87B02-GAL4 crossed to MCFO-7 |
| S2G | VT016795-GAL4 crossed to MCFO-7 |
| S2H | VT016279-GAL4 crossed to MCFO-7 |

**Movie S1.** (MovieS1.avi)

**Rotating movie of the VS, T4d, T5d, and LPi3-4 cells. Related to Figure 3.**

VS: magenta, VS dendrite: green, T4d: shades of green, T5d: shades of blue, LPi3-4: red and orange, presynapse: yellow, postsynapse: white.

**File S1. Connections of the T4 and T5 cells. Related to Figure 2A.**

**S1A**: List of input neurons and numbers of synapses of the core T4 and T5 neurons (20 cells each)

**S1B**: List of output neurons and numbers of synapses of the core T4 and T5 neurons (20 cells each)

**File S2. Outputs of the HS and VS branches. Related to Figure 3F.**

**S2A**: List of input neurons and numbers of synapses of HS cell branches

**S2B**: List of input neurons and numbers of synapses of VS cell branches in Lop2

**S2C**: List of input neurons and numbers of synapses of VS cell branches in Lop4

**File S3. Connections of the bilayer LPi cells. Related to Figure 4.**

**S3A**: List of input neurons and numbers of synapses of an LPi1-2 cell branch

**S3B**: List of output neurons and numbers of synapses of an LPi1-2 cell branch

**S3C**: List of input neurons and numbers of synapses of an LPi2-1 cell branch

S3D: List of output neurons and numbers of synapses of an LPi2-1 cell branch

**S3E**: List of input neurons and numbers of synapses of an LPi3-4 cell branch

**S3F**: List of output neurons and numbers of synapses of an LPi3-4 cell branch

**S3G**: List of input neurons and numbers of synapses of an LPi4-3 cell branch

**S3H**: List of output neurons and numbers of synapses of an LPi4-3 cell branch

## Notes

http://emdata.janelia.org/optic-lobe/

### Competing Interest Statement

The authors have declared no competing interest.

## References

1. Borst, A., Haag, J., and Reiff, D.F. (2010). Fly Motion Vision. Annual Review of Neuroscience 33, 49–70. 10.1146/annurev-neuro-060909-153155.

2. Ache, J.M., Polsky, J., Alghailani, S., Parekh, R., Breads, P., Peek, M.Y., Bock, D.D., von Reyn, C.R., and Card, G.M. (2019). Neural Basis for Looming Size and Velocity Encoding in the *Drosophila* Giant Fiber Escape Pathway. Current Biology 29, 1073–1081.e1074. https://doi.org/10.1016/j.cub.2019.01.079.

3. Klapoetke, N.C., Nern, A., Peek, M.Y., Rogers, E.M., Breads, P., Rubin, G.M., Reiser, M.B., and Card, G.M. (2017). Ultra-selective looming detection from radial motion opponency. Nature 551, 237–241. 10.1038/nature24626.

4. Morimoto, M.M., Nern, A., Zhao, A., Rogers, E.M., Wong, A.M., Isaacson, M.D., Bock, D.D., Rubin, G.M., and Reiser, M.B. (2020). Spatial readout of visual looming in the central brain of Drosophila. Elife 9. 10.7554/eLife.57685.

5. Song, B.M., and Lee, C.H. (2018). Toward a Mechanistic Understanding of Color Vision in Insects. Front Neural Circuits 12, 16. 10.3389/fncir.2018.00016.

6. Schnaitmann, C., Haikala, V., Abraham, E., Oberhauser, V., Thestrup, T., Griesbeck, O., and Reiff, D.F. (2018). Color Processing in the Early Visual System of Drosophila. Cell 172, 318–330 e318. 10.1016/j.cell.2017.12.018.

7. Behnia, R., Clark, D.A., Carter, A.G., Clandinin, T.R., and Desplan, C. (2014). Processing properties of ON and OFF pathways for *Drosophila* motion detection. Nature 512, 427–430. 10.1038/nature13427.

8. Maisak, M.S., Haag, J., Ammer, G., Serbe, E., Meier, M., Leonhardt, A., Schilling, T., Bahl, A., Rubin, G.M., Nern, A., et al. (2013). A directional tuning map of *Drosophila* elementary motion detectors. Nature 500, 212–216. 10.1038/nature12320.

9. Brand, A.H., and Perrimon, N. (1993). Targeted gene expression as a means of altering cell fates and generating dominant phenotypes. development 118, 401–415.

10. Duffy, J.B. (2002). GAL4 system in *Drosophila:* a fly geneticist’s Swiss army knife. Genesis 34, 1–15. 10.1002/gene.10150.

11. Jenett, A., Rubin, G.M., Ngo, T.T., Shepherd, D., Murphy, C., Dionne, H., Pfeiffer, B.D., Cavallaro, A., Hall, D., Jeter, J., et al. (2012). A GAL4-driver line resource for *Drosophila* neurobiology. Cell Rep 2, 991–1001. 10.1016/j.celrep.2012.09.011.

12. Simpson, J.H. (2016). Rationally subdividing the fly nervous system with versatile expression reagents. J Neurogenet 30, 185–194. 10.1080/01677063.2016.1248761.

13. Borst, A. (2009). Drosophila’s view on insect vision. Curr Biol 19, R36–47. 10.1016/j.cub.2008.11.001.

14. Fischbach, K.-F., and Dittrich, A. (1989). The optic lobe of *Drosophila melanogaster*. I. A Golgi analysis of wild-type structure. Cell and tissue research 258, 441–475.

15. Strausfeld, N.J. (1976). Atlas of an insect brain (Springer-Verlag).

16. Nern, A., Pfeiffer, B.D., and Rubin, G.M. (2015). Optimized tools for multicolor stochastic labeling reveal diverse stereotyped cell arrangements in the fly visual system. Proc Natl Acad Sci U S A 112, E2967–2976. 10.1073/pnas.1506763112.

17. Tirian, L., and Dickson, B. (2017). The VT GAL4, LexA, and split-GAL4 driver line collections for targeted expression in the *Drosophila* nervous system. bioRxiv. 10.1101/198648.

18. Rivera-Alba, M., Vitaladevuni, S.N., Mishchenko, Y., Lu, Z., Takemura, S.Y., Scheffer, L., Meinertzhagen, I.A., Chklovskii, D.B., and de Polavieja, G.G. (2011). Wiring economy and volume exclusion determine neuronal placement in the *Drosophila* brain. Curr Biol 21, 2000–2005. 10.1016/j.cub.2011.10.022.

19. Takemura, S.Y., Lu, Z., and Meinertzhagen, I.A. (2008). Synaptic circuits of the *Drosophila* optic lobe: the input terminals to the medulla. J Comp Neurol 509, 493–513. 10.1002/cne.21757.

20. Takemura, S.Y., Bharioke, A., Lu, Z., Nern, A., Vitaladevuni, S., Rivlin, P.K., Katz, W.T., Olbris, D.J., Plaza, S.M., Winston, P., et al. (2013). A visual motion detection circuit suggested by *Drosophila* connectomics. Nature 500, 175–181. 10.1038/nature12450.

21. Takemura, S.Y., Nern, A., Chklovskii, D.B., Scheffer, L.K., Rubin, G.M., and Meinertzhagen, I.A. (2017). The comprehensive connectome of a neural substrate for ‘ON’ motion detection in *Drosophila*. eLife 6, e24394. 10.7554/eLife.24394.

22. Shinomiya, K., Huang, G., Lu, Z., Parag, T., Xu, C.S., Aniceto, R., Ansari, N., Cheatham, N., Lauchie, S., Neace, E., et al. (2019). Comparisons between the ON- and OFF-edge motion pathways in the *Drosophila* brain. eLife 8, e40025. 10.7554/eLife.40025.

23. Shinomiya, K., Karuppudurai, T., Lin, T.Y., Lu, Z., Lee, C.H., and Meinertzhagen, I.A. (2014). Candidate neural substrates for off-edge motion detection in *Drosophila*. Curr Biol 24, 1062–1070. 10.1016/j.cub.2014.03.051.

24. Strother, J.A., Wu, S.T., Wong, A.M., Nern, A., Rogers, E.M., Le, J.Q., Rubin, G.M., and Reiser, M.B. (2017). The emergence of directional selectivity in the visual motion pathway of *Drosophila*. Neuron 94, 168–182 e110. 10.1016/j.neuron.2017.03.010.

25. Serbe, E., Meier, M., Leonhardt, A., and Borst, A. (2016). Comprehensive characterization of the major presynaptic elements to the *Drosophila* OFF motion detector. Neuron 89, 829–841. 10.1016/j.neuron.2016.01.006.

26. Strausfeld, N.J. (2021). The lobula plate is exclusive to insects. Arthropod Struct Dev 61, 101031. 10.1016/j.asd.2021.101031.

27. Hausen, K., and Egelhaaf, M. (1989). Neural Mechanisms of Visual Course Control in Insects. held in Berlin, Heidelberg, 1989//. D.G. Stavenga, and R.C. Hardie, eds. (Springer Berlin Heidelberg), pp. 391–424.

28. Hausen, K. (1984). The Lobula-Complex of the Fly: Structure, Function and Significance in Visual Behaviour. In Photoreception and Vision in Invertebrates, M.A. Ali, ed. (Springer US), pp. 523–559. 10.1007/978-1-4613-2743-1_15.

29. Eckert, H. (1980). Functional properties of the H1-neurone in the third optic Ganglion of the Blowfly, *Phaenicia*. Journal of comparative physiology 135, 29–39. 10.1007/BF00660179.

30. Hausen, K. (1976). Functional characterization and anatomical identification of motion sensitive neurons in the lobula plate of the blowfly *Calliphora erythrocephala*. Zeitschrift für Naturforschung c 31, 629–634.

31. Scott, E.K., Raabe, T., and Luo, L. (2002). Structure of the vertical and horizontal system neurons of the lobula plate in *Drosophila*. J Comp Neurol 454, 470–481. 10.1002/cne.10467.

32. Joesch, M., Plett, J., Borst, A., and Reiff, D.F. (2008). Response properties of motion-sensitive visual interneurons in the lobula plate of *Drosophila melanogaster*. Curr Biol 18, 368–374. 10.1016/j.cub.2008.02.022.

33. Schnell, B., Joesch, M., Forstner, F., Raghu, S.V., Otsuna, H., Ito, K., Borst, A., and Reiff, D.F. (2010). Processing of horizontal optic flow in three visual interneurons of the *Drosophila* brain. J Neurophysiol 103, 1646–1657. 10.1152/jn.00950.2009.

34. Boergens, K.M., Kapfer, C., Helmstaedter, M., Denk, W., and Borst, A. (2018). Full reconstruction of large lobula plate tangential cells in *Drosophila* from a 3D EM dataset. PLoS One 13, e0207828. 10.1371/journal.pone.0207828.

35. Mauss, A.S., Pankova, K., Arenz, A., Nern, A., Rubin, G.M., and Borst, A. (2015). Neural circuit to integrate opposing motions in the visual field. Cell 162, 351–362. 10.1016/j.cell.2015.06.035.

36. Raghu, S.V., and Borst, A. (2011). Candidate glutamatergic neurons in the visual system of *Drosophila*. PLoS One 6, e19472. 10.1371/journal.pone.0019472.

37. Mauss, A.S., Meier, M., Serbe, E., and Borst, A. (2014). Optogenetic and pharmacologic dissection of feedforward inhibition in Drosophila motion vision. J Neurosci 34, 2254–2263. 10.1523/JNEUROSCI.3938-13.2014.

38. Wu, M., Nern, A., Williamson, W.R., Morimoto, M.M., Reiser, M.B., Card, G.M., and Rubin, G.M. (2016). Visual projection neurons in the *Drosophila* lobula link feature detection to distinct behavioral programs. eLife 5. 10.7554/eLife.21022.

39. Gilbert, C., and Strausfeld, N.J. (1991). The functional organization of male-specific visual neurons in flies. J Comp Physiol A 169, 395–411. 10.1007/BF00197653.

40. Panser, K., Tirian, L., Schulze, F., Villalba, S., Jefferis, G., Buhler, K., and Straw, A.D. (2016). Automatic Segmentation of *Drosophila* Neural Compartments Using GAL4 Expression Data Reveals Novel Visual Pathways. Curr Biol 26, 1943–1954. 10.1016/j.cub.2016.05.052.

41. Davis, F.P., Nern, A., Picard, S., Reiser, M.B., Rubin, G.M., Eddy, S.R., and Henry, G.L. (2020). A genetic, genomic, and computational resource for exploring neural circuit function. Elife 9. 10.7554/eLife.50901.

42. Strausfeld, N.J., and Okamura, J.Y. (2007). Visual system of calliphorid flies: organization of optic glomeruli and their lobula complex efferents. J Comp Neurol 500, 166–188. 10.1002/cne.21196.

43. Scheffer, L.K., Xu, C.S., Januszewski, M., Lu, Z., Takemura, S.Y., Hayworth, K.J., Huang, G.B., Shinomiya, K., Maitlin-Shepard, J., Berg, S., et al. (2020). A connectome and analysis of the adult *Drosophila* central brain. eLife 9. 10.7554/eLife.57443.

44. Isaacson MD, Eliason JLM, Nern A, Rogers EM, Rubin GM, Branson K, and MB, R. (in preparation). Small-field visual projection neurons detect translational optic flow and contribute to the regulation of forward walking.

45. Shinomiya, K., Horne, J.A., McLin, S., Wiederman, M., Nern, A., Plaza, S.M., and Meinertzhagen, I.A. (2019). The Organization of the Second Optic Chiasm of the *Drosophila* Optic Lobe. Front Neural Circuits 13, 65. 10.3389/fncir.2019.00065.

46. Haag, J., Vermeulen, A., and Borst, A. (1999). The intrinsic electrophysiological characteristics of fly lobula plate tangential cells: III. Visual response properties. J Comput Neurosci 7, 213–234. 10.1023/a:1008950515719.

47. Krapp, H.G., and Hengstenberg, R. (1996). Estimation of self-motion by optic flow processing in single visual interneurons. Nature 384, 463–466. 10.1038/384463a0.

48. Hengstenberg, R. (1982). Common visual response properties of giant vertical cells in the lobula plate of the blowfly *Calliphora*. Journal of comparative physiology 149, 179–193. 10.1007/BF00619212.

49. Pierantoni, R. (1976). A look into the cock-pit of the fly. The architecture of the lobular plate. Cell Tissue Res 171, 101–122. 10.1007/BF00219703.

50. Hengstenberg, R., Hausen, K., and Hengstenberg, B. (2004). The number and structure of giant vertical cells (VS) in the lobula plate of the blowflyCalliphora erythrocephala. Journal of comparative physiology 149, 163–177.

51. Kim, A.J., Fenk, L.M., Lyu, C., and Maimon, G. (2017). Quantitative Predictions Orchestrate Visual Signaling in Drosophila. Cell 168, 280–294 e212. 10.1016/j.cell.2016.12.005.

52. Barnhart, E.L., Wang, I.E., Wei, H., Desplan, C., and Clandinin, T.R. (2018). Sequential Nonlinear Filtering of Local Motion Cues by Global Motion Circuits. Neuron 100, 229–243 e223. 10.1016/j.neuron.2018.08.022.

53. Zheng, Z., Lauritzen, J.S., Perlman, E., Robinson, C.G., Nichols, M., Milkie, D., Torrens, O., Price, J., Fisher, C.B., Sharifi, N., et al. (2018). A Complete Electron Microscopy Volume of the Brain of Adult Drosophila melanogaster. Cell 174, 730–743 e722. 10.1016/j.cell.2018.06.019.

54. Eckert, H., and Dvorak, D.R. (1983). The centrifugal horizontal cells in the lobula plate of the blowfly, *Phaenicia sericata*. Journal of Insect Physiology 29, 547–560.

55. Wei, H., Kyung, H.Y., Kim, P.J., and Desplan, C. (2020). The diversity of lobula plate tangential cells (LPTCs) in the *Drosophila* motion vision system. J Comp Physiol A Neuroethol Sens Neural Behav Physiol 206, 139–148. 10.1007/s00359-019-01380-y.

56. Gauck, V., Egelhaaf, M., and Borst, A. (1997). Synapse distribution on VCH, an inhibitory, motion-sensitive interneuron in the fly visual system. J Comp Neurol 381, 489–499.

57. Meyer, E.P., Matute, C., Streit, P., and Nassel, D.R. (1986). Insect optic lobe neurons identifiable with monoclonal antibodies to GABA. Histochemistry 84, 207–216. 10.1007/BF00495784.

58. Haag, J., and Borst, A. (2001). Recurrent network interactions underlying flow-field selectivity of visual interneurons. J Neurosci 21, 5685–5692.

59. Farrow, K., Haag, J., and Borst, A. (2003). Input organization of multifunctional motion-sensitive neurons in the blowfly. J Neurosci 23, 9805–9811.

60. Duistermars, B.J., Care, R.A., and Frye, M.A. (2012). Binocular interactions underlying the classic optomotor responses of flying flies. Front Behav Neurosci 6, 6. 10.3389/fnbeh.2012.00006.

61. Farrow, K., Haag, J., and Borst, A. (2006). Nonlinear, binocular interactions underlying flow field selectivity of a motion-sensitive neuron. Nat Neurosci 9, 1312–1320. 10.1038/nn1769.

62. Bishop, C.A., and Bishop, L.G. (1981). Vertical motion detectors and their synaptic relations in the third optic lobe of the fly. J Neurobiol 12, 281–296. 10.1002/neu.480120308.

63. Dvorak, D.R., Bishop, L.G., and Eckert, H.E. (1975). On the identification of movement detectors in the fly optic lobe. Journal of comparative physiology 100, 5–23.

64. Takemura, S.-y., Xu, C.S., Lu, Z., Rivlin, P.K., Parag, T., Olbris, D.J., Plaza, S., Zhao, T., Katz, W.T., Umayam, L., et al. (2015). Synaptic circuits and their variations within different columns in the visual system of *Drosophila*. Proceedings of the National Academy of Sciences 112, 13711–13716. 10.1073/pnas.1509820112.

65. Clements, J., Dolafi, T., Umayam, L., Neubarth, N.L., Berg, S., Scheffer, L.K., and Plaza, S.M. (2020). neuPrint: Analysis Tools for EM Connectomics. bioRxiv, 2020.2001.2016.909465. 10.1101/2020.01.16.909465.

66. Feng, L., Zhao, T., and Kim, J. (2015). neuTube 1.0: a new design for efficient neuron reconstruction software based on the SWC format. eneuro 2, ENEURO.0049-0014.2014.

67. Otsuna, H., and Ito, K. (2006). Systematic analysis of the visual projection neurons of *Drosophila melanogaster*. I. Lobula-specific pathways. J Comp Neurol 497, 928–958. 10.1002/cne.21015.

